# The emergence of integrated information, complexity, and consciousness at criticality

**DOI:** 10.1101/521567

**Authors:** Sina Khajehabdollahi, Pubuditha M. Abeyasinghe, Adrian M. Owen, Andrea Soddu

## Abstract

Using the critical Ising model of the brain, integrated information as a measure of consciousness is measured in toy models of generic neural networks. Monte Carlo simulations are run on 159 random weighted networks analogous to small 5-node neural network motifs. The integrated information generated by this sample of small Ising models is measured across the model parameter space. It is observed that integrated information, as a type of order parameter not unlike a concept like magnetism, undergoes a phase transition at the critical point in the model. This critical point is demarcated by the peaks of the generalized susceptibility of integrated information, a point where the ‘consciousness’ of the system is maximally susceptible to perturbations and on the boundary between an ordered and disordered form. This study adds further evidence to support that the emergence of consciousness coincides with the more universal patterns of self-organized criticality, evolution, the emergence of complexity, and the integration of complex systems.

**Author summary:** Understanding consciousness through a scientific and mathematical language is slowly coming into reach and so testing and grounding these emerging ideas onto empirical observations and known systems is a first step to properly framing this ancient problem. This paper in particular explores the Integrated Information Theory of Consciousness framed within the physics of the Ising model to understand how and when consciousness, or integrated information, can arise in simple dynamical systems. The emergence of consciousness is treated like the emergence of other classical macroscopic observables in physics such as magnetism and understood as a dynamical phase of matter. Our findings show that the sensitivity of consciousness in a complex system is maximized when the system is undergoing a phase transition, also known as a critical point. This result, combined with a body of evidence highlighting the privelaged state of critical systems suggests that, like many other complex phenomenon, consciousness may simply follow from/emerge out of the tendency of a system to self-organize to criticality.

## Introduction

A growing body of evidence in the past few decades has emerged suggesting that many disparate natural and particularly biological phenomena reside in a critical regime of dynamics on the cusp between order and disorder [1–10]. This seemingly ubiquitous phenomena has sparked a renaissance of new ideas attempting to understand the self-organizing nature of our world [11]. More specifically, it has been shown that critical models, and in particular the Ising model at criticality, model the statistics of brain dynamics quite well [12–15], which combined with evidence of critical variables in brain dynamics has led to the emergence of the critical brain hypothesis [8,10]. Systems tuned to criticality, self-organized or otherwise, exhibit a number of useful informational properties that allow for the efficient distribution of and susceptibility to information [6,10,15–17]. These ideas have been further developed to suggest more broadly that critical systems are evolutionary advantageous and stable attractors for systems living in complex environments as they are more effective at reacting to their environment and ensuring their continued survival [18–20].

### Integrated Information Theory

In order to approach the problem of consciousness a working definition is needed and as such, this project attempts to understand and explore one emerging model known as integrated information theory (IIT) [21]. IIT is a top-down, phenomenological approach to defining consciousness [21]. Starting from phenomenological axioms the theory constructs mathematical postulates that create a workspace for researchers to test and explore this particular definition of consciousness and all its associated controversies. Ultimately, the main measure proposed by IIT is the mathematical object called integrated information (Φ) (Big Phi) which generally seeks to measure ‘how much the whole is greater than the sum of its parts’ of a causal structure. Though other measures exist [22] which try to capture some form of integration or complexity, this paper will use Φ as its main metric. For a wholesome overview of the mathematical taxonomy of the possible variations in defining integrated information, see [23].

Unfortunately, many calculations in the theory prove to be intractable, scaling super-exponentially with respect to the size of the system of interest, resembling a high-dimensional ‘traveling salesman’ problem at certain stages of the algorithm. To measure integrated information one needs to have access to the transition probabilities of the system. In other words, something like the partition function, the Markov matrix, or the structure of the causal model is needed. Naturally this is information we are not always privy to when it comes to complex phenomena like brain dynamics. This problem is circumvented by using a sufficiently simple model where the transition rates can be readily calculated. Furthermore, if one wants to understand the human brain and experience of consciousness within the language of IIT, some sort of bridge needs to be built to link IIT and brain dynamics. Therefore, to solve the problem of computational intractability and contextualize our work within the human condition, the Ising model of the brain is employed to address both problems.

### The Critical, Generalized Ising Model

The generalized Ising model acts as this bridge by virtue of the model’s ability to exhibit phase transitions and critical points as well as being the simplest (max entropy) model associated with empirical pairwise correlation data [18,24]. Historically, it was the 2D Ising model that first demonstrated the ability to exhibit a phase transition at some critical temperature T_c_, a global scaling parameter of the model. More recently, it has been shown to also exhibit similar statistical structures to that of the brain which is also thought to be a critical system and has given rise to the Critical Brain Hypothesis [2–10,14,16,25–29]. This quality allows this seemingly crude and simple model to relatively accurately recreate the pairwise correlation structures and distributions to that of the human brain and acts as a proxy to the brain simply by sharing some universal characteristic with it. By taking advantage of this, albeit roughly defined, shared universality class between the critical Ising model and the Brain, we can study the Ising model through the scope of IIT and attempt to project our results onto the Brain.

The crudeness of the 2D Ising model can be slightly overcome by generalizing its interactions such that they are not confined to only nearest-neighbors and instead can use any general structural connectivity that is given as input. This generalization allows us to model not just 2D lattice interactions but also more generally, simple neural network motifs. To this end, the Ising model is simulated on 159 randomly generated, positive weighted networks to explore the combinatoric space of neural network motifs. Each unique network exhibits its own idiosyncratic phase transition as measured across a variety of its natural variables. Φ and its susceptibility *χ*_Φ_ is also measured across the parameter space of these models, fitting in quite naturally among the other natural variables commonly used in the physical paradigm such as Energy (E), Magnetization (M), or Magnetic Susceptibility (χ). Simulations sweep across the model’s only parameter, the temperature of its surrounding heat bath. As this parameter is swept from low to high temperatures, larger and larger energetic fluctuations become likely. In many cases, as this parameter is swept the organizational structure of the system can dramatically change, exhibiting a phase transition. This abrupt change in quality as a result in a continuous change in a particular quantity is at the heart of many of the most interesting complex systems such as genetic networks [30] societal organizations, financial markets [31,32], or swarming behaviours [33,34]. The critical points where these transitions are located are demarcated by the ‘critical temperature’ (*T_c_*) of our generalized model.

### Schematic Overview

In Equation [1] an overview of the strategy employed in this paper is summarized. Ising simulations are run given a connectivity matrix and temperature (**J**, *T*) with the outputs: correlation matrix, magnetization, integrated information, and the susceptibilities of magnetization and integrated information, all as a function of the temperature [***ρ***^sim^, *M*, Φ, *χ, χ*_Φ_] (*T*). These outputs are used to identify the critical temperature of the model as well as allow the observation of the behaviour of integrated information in the model.

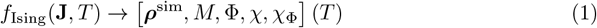

### Emergence of Complexity

Results indicate that the integrated information generated in the Ising model, much like the classical variable ‘Magnetization’ as a macroscopic order parameter, undergoes a phase transition at the model’s critical point. This is detected by locating the peaks of its susceptibility curves as a function of temperature [35]. This result is important because it indicates that the integrated information structure of simple neural networks can behave critically, exhibiting maximal susceptibility to perturbations and allowing for a form of consciousness that balances coherence and continuity with information and variance. If integrated information is a good description of consciousness, this implies that the phenomenon of consciousness may just be the next branch in the big family tree of phase transitions and an evolutionary attractor. These results fit into a larger paradigm that seeks to understand the nature of evolution and the adaptive advantage of critical systems in the context of a universe undergoing a cascade of phase transitions [36,37].

## Results

### Model Simulations

159 Ising simulations were generated using N = 5 nodes, fully-connected networks with random weights. Summary statistics were calculated for all simulations as a function of the fitting parameter *T*: magnetization *M*, integrated information Φ, the magnetic susceptibility *χ*, the generalized susceptibility of integrated information *χ*_Φ_, and variances across all the random network samples 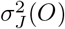.

Averaging these variables across all random networks shows a strong parallel between the behaviour of Magnetization M and integrated information Φ generated by the system (Fig 1). Near the onset of criticality (generally be approximated by the peak of the magnetic susceptibility curve [35]) integrated information, much like the magnetization in the Ising model, undergoes a phase transition which is seen as a peak in the susceptibility of Φ. The regime where the fluctuations of integrated information are maximized suggests a transition point for integrated information as an order parameter. Our results predict that the phenomenon of criticality extends into the behaviour of integrated information on the Ising model.

**Fig 1.**
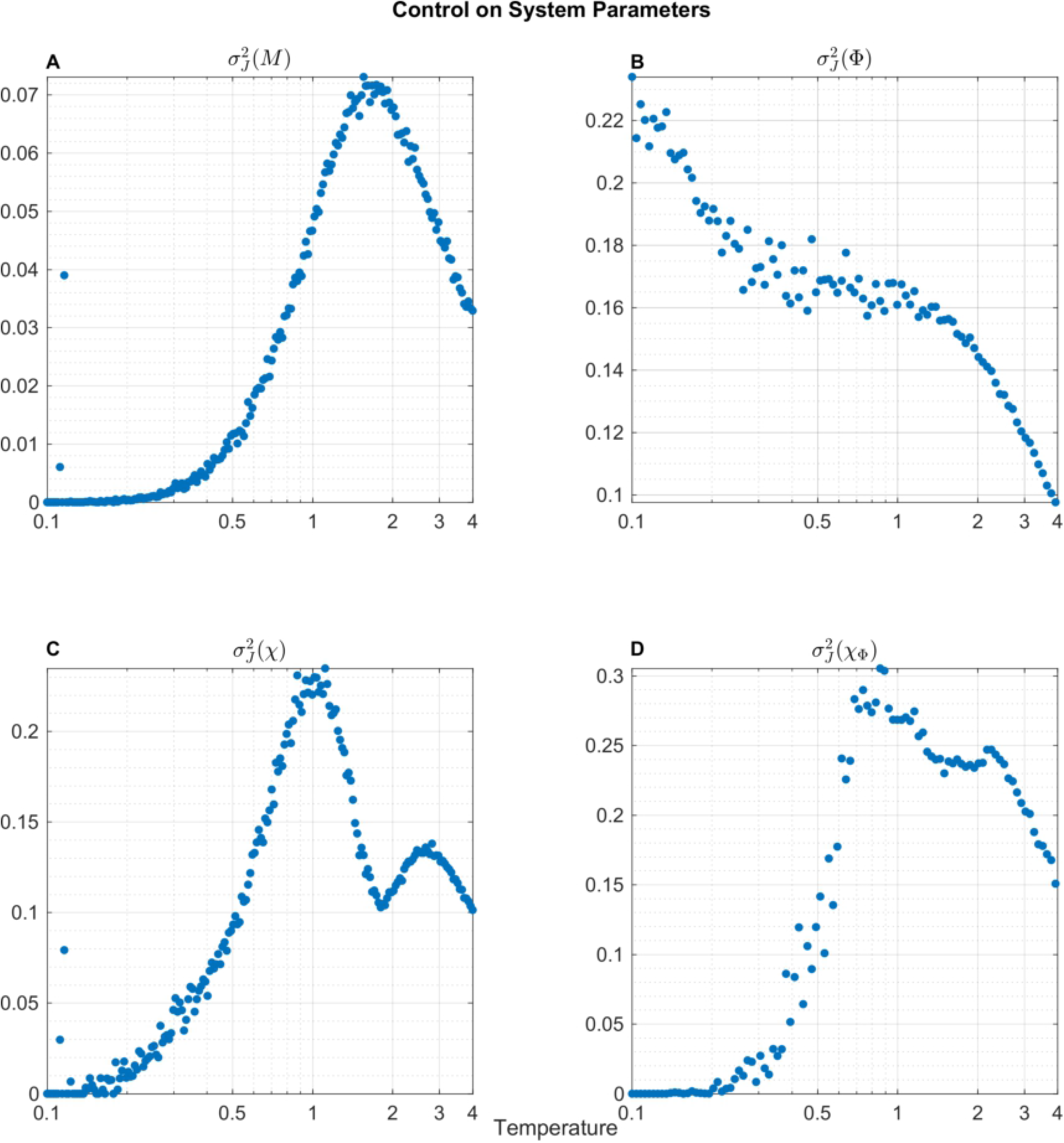
The summary statistics for the two order parameters, Magnetization *M* and Φ (panels **A, B**) across all the 159 random network simulations are shown. The variance of Φ, 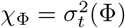 (panel **D**) is interpreted as a susceptibility of Φ and is compared to the magnetic susceptibility *χ* (panel **C**). These susceptibilities peak at the same critical temperature indicating the phase transition of integrated information as an order parameter in the Ising model.

Of the 159 of random networks simulated, only 6 demonstrated the ability to maximize Φ at criticality while the rest had the general tendency to decrease Φ as a function of temperature, though not necessarily monotonically. A peculiar behaviour observed in a symmetry breaking of integrated information is that individual Φ(*T*) functions are split into two branches near the critical point. This peculiarity is still not yet understood as the causal structure of the Ising model is expected to be symmetric under system flips.

In Fig 1 summary statistics for the order and susceptibility parameters of the random networks are shown. Magnetization *M* and its corresponding susceptibility *χ*, are plotted in the left-most column, and Φ and its susceptibility *χ*_Φ_ are plotted in the second column. In Fig 2 the variance of these variables across the different random network connectivities 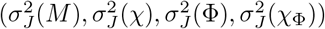 are shown. The variances 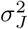 illustrate the control exhibited by the choice of connectivity onto the order parameters across the uniformly sampled random networks, whereas the susceptibilities quantify the mean fluctuations of those order parameters within each random network, averaged across all random networks. These summary statistics give first-order insights into the diagnosis and control of simple neural networks. We note that at the critical temperature, denoted roughly by the peaks of *χ*, the susceptibility of Φ, *χ*_Φ_ also peaks. When looking at Φ across different simulations, 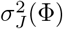, we observe that there seems to be two transition points. One transition point at low temperatures leading into a plateau region followed by a second transition close to the classical critical point where the variations in Φ begin to fall off. These results highlight the regions where changes in the structural connectivity of the model have the most influence on the generation of integrated information. While the magnetization of the model near criticality is maximally sensitive to changes in the structural connectivity, integrated information instead has a broad plateau region of uniform sensitivity. This result is useful in assessing how structural changes in a system can lead to functional changes which are capable of generating integrated information or consciousness.

**Fig 2.**
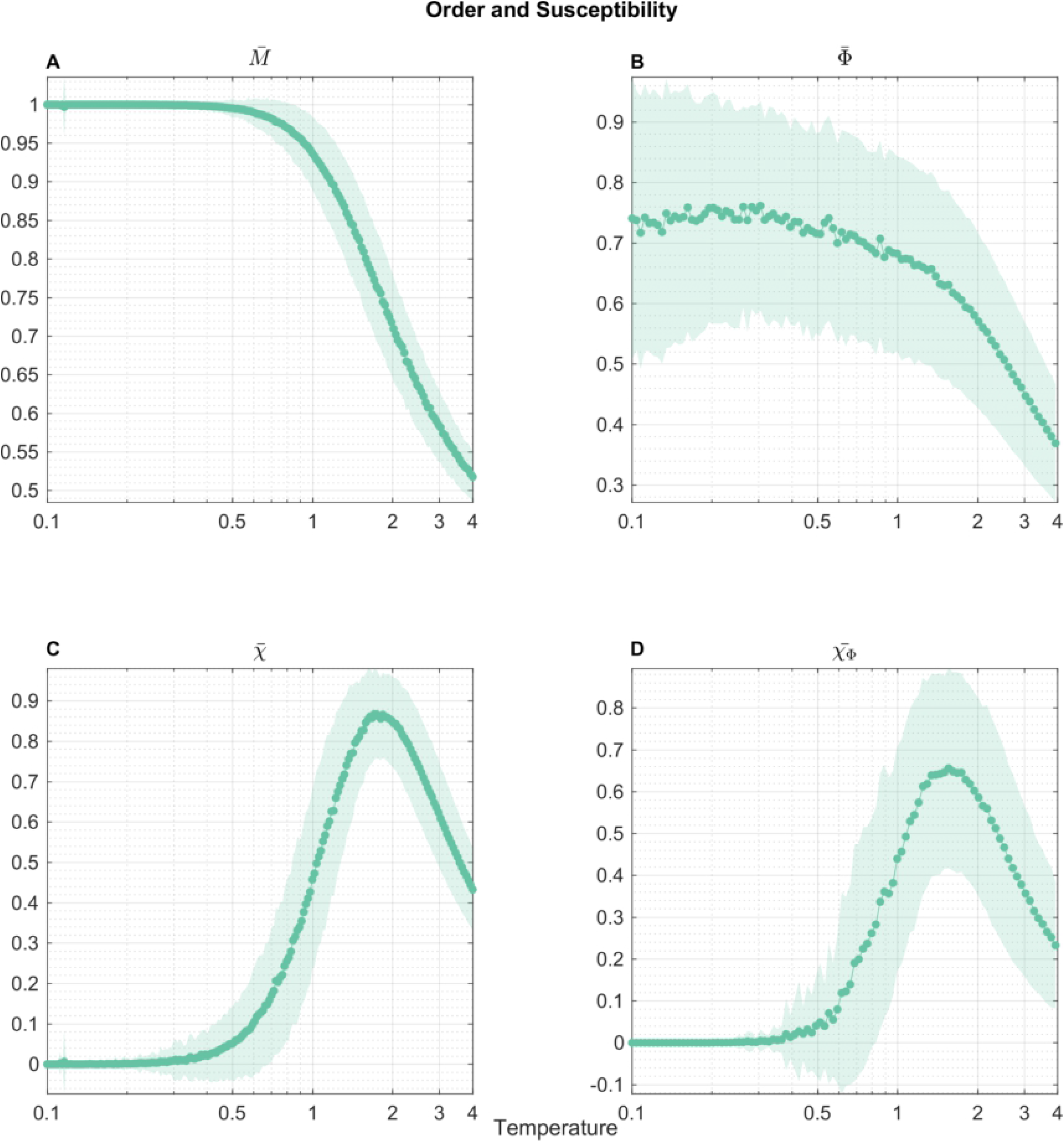
The variance of the order parameters *M*, Φ (panels **A, B**) and their susceptibilities (panels **C, D**) across different connectivities are plotted. These plots demonstrate the potential control one can impart on the Ising model by changing the connectivity matrix. Notably, the potential control reaches a local minima at the critical temperatures of the model, indicating a convergence to a universality class. However, local maxima surrounding the critical point indicate the possibility to control the transitions towards criticality.

## Discussion

### Phase Transitions & IIT

To investigate the properties of this new measure of integrated information introduced by IIT we have in this study employed the relatively simple Ising model to act as a proxy to the real brain. The Ising model is generalized to use any graph as its connectivity where in this study we have looked at 159 random networks of 5 nodes. The results from the Ising model analyzed with IIT show that integrated information tends to be maximally susceptible at the critical temperature (Fig 1). The statistics of the 159 random networks summarize these results across variations of fully connected connectivity matrices to show that while there exists a rich variety of Φ(*T*) curves, on average the ‘susceptibility’ of Φ(*T*), (*χ*_Φ_(*T*)), behaves quite similarly to the magnetic susceptibility that is normally the marker for the second order phase transition of the classical 2D Ising model. These results indicate that integrated information can more broadly be considered as a macroscopic order parameter which seems to undergoes a phase transition at the critical temperature of the model. To generate a taxonomy of the possible phases that integrated information could exhibit would require a much more thorough exploration of the possible structural connectivities and dynamical rules that a system could obey. This project confined itself to the Ising model on fully connected graphs obeying the Metropolis algorithm. In the future as more efficient algorithms for calculating Φ emerge (or as a compromise accurate correlates of Φ) combined with Monte Carlo and network renormalization group methods [16,40–45] the exploration of larger networks of different classes (e.g. sparse, modular hierarchical, small-world, fractal) could lead to the identification of a rich taxonomy of phases of integrated information.

### Control

When considering the control that can be exerted onto this Ising model by changing network edge weights, it is clear that *at* the critical point, the system reaches a local minima in the influence that changing the connectivity of the network can impart on the system’s general susceptibilities (Fig 2, panels **C, D**). This is consistent with the notion that these critical Ising regimes converge to identical (or similar) universality classes and therefore varying the networks in this regime imparts a minimal change in the general susceptibility of the system. However, on the edge of criticality, the generalized susceptibilities’ exhibit local maxima, both in the case of the magnetic susceptibility *χ* and the integrated information susceptibility *χ*_Φ_. In other words, on the *approach* towards criticality, maximum control can be exerted onto such networks when it comes to modulating their general susceptibilities. In fact, there is some evidence to believe that the brain is not *exactly* critical and rather it deviates slightly towards sub-criticality [46,47]. Therefore this experiment gives credence to the notion that a *sub-critical* system may be easier to control and therefore lends evolutionary/adaptive advantages in life. This control only begins to decrease in the super-critical regime where other properties of the system also begin to fall apart and any evolutionary advantage is likely lost.

### Evolution & Complexity

The exploration of integrated information in the context of critical systems undergoing phase transitions motivates a few new questions in regards to the relationship between evolution, complexity, and consciousness. In the work done by [55, 56] on complexity and the evolution of neural models and integrated information, it was shown that fitness can correlate strongly with Φ when the system is constrained in size/resources. While it is not always true that a system will evolve to generate high Φ under more liberal constraints (infinite resources), it does seem to be that there may be some evolutionary advantage for having high Φ. Since Φ essentially measures the emergence of higher-order concepts within a system, intuitively it may not be surprising that systems that are capable of generating higher-order concepts will be capable of representing and reacting to a more diverse set of states than systems that cannot. Therefore for resource-limited systems, having an efficient means to represent internal and external states may automatically give rise to high Φ or consciousness.

It is fair to think of integrated information as a type of complexity measure as it aims to measure how mechanisms in a system interact and constrain each other in emergent and irreducible ways. The theory aims to measure emergent properties of a system that cannot be explained by independent (or semi-independent) components of that system. The measure is sensitive to not just information, which in general can be maximized by deterministic systems with unique pasts and futures, but also to the distribution and integration of information which in general can be maximized by strongly coupled systems. To have a system that is both strongly coupled and informative requires a balance between segregating forces that act to differentiate the system into diverse states as well as coherent, integrating forces that create new forms of information that could not otherwise arise from the individual components. In a system like the Ising model, it is expected that these exact properties emerge near the critical temperature at the onset of its phase transition.

### Utility of Criticality

By definition, critical systems have diverging correlation lengths, undergo critical slowing-down (integration in space and time), and simultaneously exhibit distinct and segregated structures at all scales (scale-invariance). They are generally found in regimes of systems undergoing some kind of transition between different phases (e.g. magnetized vs. non-magnetized in the Ising model, or synchrony vs. asynchrony in the Kuramuto model [48–52]). In contrast to sub-critical regimes which can become completely uniform due to their strong coupling (high integration, low differentiation) and super-critical regimes which can become completely noise driven (low integration, high differentiation), critical systems sit in the sweet spot to generate non-negligible Φ that is maximally susceptible to the perturbations of its environment and its own state. Our results indicate that while sub-critical regimes are capable of generating high Φ, the variations in Φ in this regime are negligible. Only near the critical point does Φ have both large values and large fluctuations indicating that the critical point of the system is maximally receptive and responsive to its own states.

### Conclusion

The novelty of our study is best framed in the larger context of the emerging complexity of our world [36] and criticality in physics where the definitions suggested by [53,54] go so far to define complexity as the ‘‘tendency of large dynamical systems to organize themselves into a critical state”. For example it has been shown that in neural tissues and in a cortical branching model (which is not too different from the classical Ising model) that neural complexity is maximized at criticality [6] or that the minimal complexity of adapting agents increases with fitness [55]. Even more broadly it has been shown that criticality may be useful for learning [9], and for optimizing information processing [10,18,19]. Therefore, the novelty of this study is that by using the framework of IIT to define consciousness, we show evidence that consciousness as a property of matter can undergo a phase transition at criticality which, combined with previous evidence that the brain may be critical, suggests that consciousness as we know it may simply arise out of the tendancy of the brain to self-organize towards criticality. The importance of this conclusion lies in the fact that this ancient problem of understanding and defining consciousness may ultimately be best framed and understood within the physics and evolutionary dynamics of self-organizing critical systems.

### Future Work

It is clear that both the magnitude and susceptibility of Φ in the Ising model (and in general) are extremely sensitive on the exact nature of its connectivity and the dynamic rules that govern it. The nature of a phase transition that a system may exhibit is also contingent on these properties (though disparate systems can fall into the same universality classes). So far our simulations were run on static networks, but in general one can let the network itself evolve. Future work interested in how the networks themselves arise could explore different evolution algorithms under different dynamical rules in combination with the analysis from IIT to assess the role of evolution and environment in generating Φ. Exploring the behaviour of Φ in different classes of phase transitions could further develop the ideas behind the critical brain hypothesis and coerce the fields of neuroscience, complexity science, material science, and statistical mechanics to work together to understand the brain. In a way that is analogous to the modular but integrative organization of the brain, these distinct disciplines in science will need to integrate with each other if humanity hopes to understand complex integrative systems like the brain, the societies we live in, and the cultural, or economical and socio-political organizations that have emerged with the rise of human civilization.

## Materials and methods

### Ising Model

The Ising model is one of the simplest ways to model many-body interactions between simple elements. Traditionally these elements are described as ‘spins’ which can be in one of two states, *s_i_* = {±1}. Though the implementation of the model started from the humble origins of modeling the macroscopically observed phenomenon of phase transitions, the Ising model has found applications and generated insights in almost all domains of life. Since the ‘spins’ in the model are abstractions, the ‘spins’ can represent any element that can be described in a binary state. When modeling the brain, these spin elements represent neurons or clusters of neurons with ‘firing’ or ‘not firing’ binary states. Furthermore, the model in its original 2D format was organized into a lattice grid with spin elements interacting only with their nearest-neighbours and a heat bath of temperature *T*. In the generalized Ising model, we no longer constrain the system to be in a 2D lattice and instead allow any general graph to describe the connectivity *J_ij_* between elements *i* and *j*. The energy of the model in a particular configuration *s* is given by:

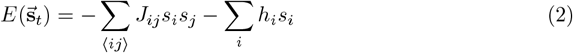

where the summation 〈*ij*〉 is over all connected elements and *h_i_* is a locally applied field or bias (which is set to 0 in this experiment). In order to bring the model to equilibrium, Metropolis update rules are used where a random element in the model is chosen and allowed the possibility for a ‘spin-flip’. A spin-flip will occur if the energy of the system decreases after flipping, or if the energy increases, then the flip will occur with a probability given by the Boltzmann factor:

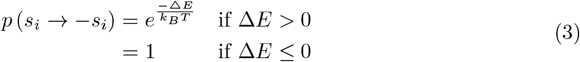

The temperature in the model affects the rate at which ‘unfavorable’ spin-flips occur; increasing the temperature increases the noise/randomness of the model’s dynamics. Within each time step in the model, all spins have the opportunity to flip once, updating simultaneously for the next step until the process is repeated for some desired number of times steps. When the system has had enough time to equilibriate past its transient initial state, observables in the model are measured repeatedly and accumulated to generate equilibrium expectation values.

### Summary Statistics

The summary statistics for observables 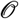 measured in this experiment are defined below. The magnetic susceptibility is shown separately.

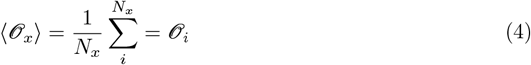

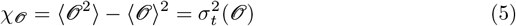

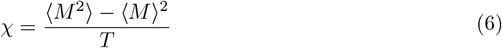

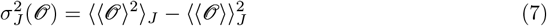

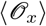 is the expectation value of an observable across some dimension *x* and for *x* = *t* time steps, *N_t_* = 2000 is the number of observations in a simulation after an initial 500 time steps to thermalize the system. 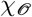 is the generalized susceptibility [19,38,39] and 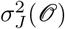 is the variance of an observable across all networks. The magnetic susceptibility (*χ*) is written separately as it is derived from the Free Energy potentials of the Ising model.

### Random Networks

159 fully connected networks of 5 nodes with random edge weights uniformly sampled between 0 and 1 are generated. The networks are then normalized such that their strongest weight is always unity. These random networks are saved as connectivity matrices and fed into the Monte Carlo Ising simulations. These random networks are designed to explore the combinatoric space of the neural network motifs possible when constrained by 5 nodes.

### Phi

Integrated Information (Φ) is calculated (using the pyPhi python toolbox [21]) in the 5-node Ising model for 2000 iterations after the model reaches a steady-state which is assumed to be achieved after 500 iterations. To calculate Φ a transition probability matrix (TPM) must be supplied which we can denote as a Markov matrix (M), as well as the configuration of the system at that time-step 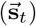 (Equation 8). Calculating the TPM requires the calculation of probablities from any configuration to any other by iterating Equation 3 across all spin sites that need to flip for the transition to occur and taking their product.

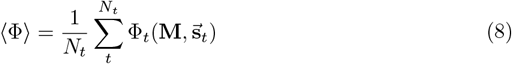

## Acknowledgments

We thank Dr. Larissa Albantakis for her detailed correspondence on ITT which greatly accelerated the progress of this project. Similarly, we thank Will Mayner for his detailed correspondence and assistance in troubleshooting the IIT pyPhi toolbox and getting simulations up and running. Adrian M. Owen is supported by a Canada Excellence Research Chair (CERC) Award (#215063).

